# Contribution of host species and pathogen clade to snake fungal disease hotspots in Europe

**DOI:** 10.1101/2022.11.10.515990

**Authors:** Gaëlle Blanvillain, Jeffrey M. Lorch, Nicolas Joudrier, Stanislaw Bury, Thibault Cuenot, Michael Franzen, Fernando Martínez-Freiría, Gaëtan Guiller, Bálint Halpern, Aleksandra Kolanek, Katarzyna Kurek, Olivier Lourdais, Alix Michon, Radka Musilová, Silke Schweiger, Barbara Szulc, Sylvain Ursenbacher, Oleksandr Zinenko, Joseph R. Hoyt

## Abstract

1. Infectious diseases are influenced by interactions between host and pathogen, and are rarely homogenous across the landscape. Areas with elevated pathogen prevalence maintain a high force of infection, can facilitate pathogen spread to new regions, and may indicate areas with impacts on host populations. However, isolating the ecological processes that result in increases in infection prevalence and intensity remains a challenge.

2. Here we elucidate the contribution of pathogen clade and host species in disease hotspots of *Ophidiomyces ophidiicola,* the pathogen that causes snake fungal disease, in 21 species of snakes infected with multiple pathogen strains across 10 countries in Europe.

3. We found isolated areas of disease hotspots in a landscape where infections were otherwise low. *O. ophidiicola* clade had important effects on transmission, and areas with multiple pathogen clades had higher host infection prevalence. Snake species identity further influenced infection, with most positive detections coming from the *Natrix* genus. Most species present in the community only experienced increased levels of infection when multiple strains were present. However, one species, *N. tessellata*, appeared highly susceptible, having increased infection prevalence regardless of pathogen strain, indicating that this species may be important in pathogen maintenance.

4. Our results suggest that both host and pathogen identity are essential components contributing to increased pathogen prevalence. More broadly, our findings indicate that coevolutionary relationships between hosts and pathogens may be key mechanisms explaining variation in landscape patterns of disease.

## Introduction

Infectious diseases can shape ecological communities by altering host abundance and distributions across the landscape (LaDeau et al. 2007, Holdo et al. 2009, Langwig et al. 2012). Disease outcomes are determined by host-pathogen interactions which are multifaceted and can interact with environmental conditions, creating a mosaic of disease hotspots across broad spatial scales (Krauss et al. 2010, Paull et al. 2012, Brown et al. 2013, Wilber et al. 2020). Hotspots of high pathogen prevalence may represent potential areas of continued impacts to host populations, serve as a source for pathogen dispersal, and maintain high propagule pressure within host communities (Kilpatrick et al. 2006, Krauss et al. 2010, Paull et al. 2012, Brown et al. 2013, Wilber et al. 2020, Laggan et al. 2022).

Heterogeneity in innate species susceptibility is recognized as a strong force influencing pathogen transmission and disease impacts for multi-host pathogens (van Riper et al. 1986, LaDeau et al. 2007, Voyles et al. 2009, Langwig et al. 2017). The distribution of highly susceptible species can determine areas of high prevalence if they are critical in pathogen maintenance (Haydon et al. 2002, Ashford 2003). However, the disproportionate contribution of a particular species may be modified by differences in community structure, environmental conditions among patches, and variation in pathogen virulence (Balaz et al. 2014, Wilber et al. 2020). Pathogen replication rates can also differ among strains and across the landscape, producing additional variation in disease prevalence (O’Hanlon et al. 2018, Greener et al. 2020). Pathogen strains with high growth rates and virulence may be due to multiple factors, including the introduction of novel strains to new locations or hosts (Li et al. 2004, Becker et al. 2017), ease or independence of transmission from affected hosts (Hawley et al. 2013, Pandey et al. 2022), and the development of novel mutations or adaptations that facilitate the escape from host resistance (McLeod and Gandon 2022). Although the interaction between host species and pathogen identity is rarely examined, theory suggests that presence of highly resistant host species could modify the effects of pathogens with high replication rates, creating cold spots of transmission across the landscape (Gandon and Michalakis 2000). Conversely, highly transmissible and virulent pathogen strains in the presence of more moderately affected host species could drive hotspots of infection (Urbina et al. 2018, Ribeiro et al. 2019, McClure et al. 2020).

The fungal pathogen *Ophidiomyces ophidiicola,* that causes snake fungal disease (SFD, also called ophidiomycosis), has been documented in over 42 species of wild snakes across three continents (Lorch et al. 2016, Burbrink et al. 2017, Franklinos et al. 2017, Meier et al. 2018, Allender et al. 2020, Davy et al. 2021, Grioni et al. 2021, Sun et al. 2021), and is considered a serious threat to the conservation of snake populations (Sutherland et al. 2014, Allender et al. 2015). Clinical signs of disease caused by *O. ophidiicola* can range from mild skin lesions, from which snakes can recover, to severe infections that impair movement, disrupt feeding behavior, and can ultimately lead to death (Lorch et al. 2016). Although population declines associated with SFD have been documented in some species of North American snakes (Lorch et al. 2016), no such declines have been reported in species across Europe, where *O. ophidiicola* has most likely coexisted with snakes for longer periods of time (Ladner et al. 2022). However, only limited information is available on *O. ophidiicola* infections across Europe, with just a few individual snakes confirmed to be infected with this pathogen from mainland Europe (Franklinos et al. 2017, Marini et al. 2023, Meier et al. 2018, Origgi et al. 2022).

Little is known of the origin of *O. ophidiicola*, and to date, three distinct clades of *O. ophidiicola* have been described: clade I, which has been found exclusively in wild snakes in Europe; clade II, which has been reported in wild snakes in North America and Taiwan as well as in captive snakes on multiple continents; and clade III, which has only been found in captive snakes (Ladner et al. 2022). The estimation of the most recent common ancestor between clade I and II (around 2000 years ago), as well as a lack of nonrecombinant intermediates in North America strongly indicate that *O. ophidiicola* was introduced to North America, potentially through multiple introduction events (Ladner et al. 2022). More recently, genotyping from a limited number of samples indicated the presence of both clades I and II in Switzerland dating back to at least 1959 (Origgi et al. 2022). In addition, slower growth rates have been reported for clade I strains of *O. ophidiicola*, suggesting that they may be less virulent than clade II strains (Franklinos et al. 2017). However, prevalence and disease severity associated with the two strains and how they influence landscape patterns of disease has not been directly compared.

To investigate the macroecological patterns of SFD across Europe we examined host and pathogen factors that may influence pathogen prevalence and disease severity across the landscape. We evaluated the presence of pathogen hotspots across Europe, examined differences in pathogen prevalence and disease severity among host species, and determined the effect of *O. ophidiicola* clade on pathogen prevalence and lesion severity. Finally, we combined host species and pathogen clade to explore factors that may contribute to areas with high pathogen prevalence across the landscape.

## 5. Methods

### (a) Location and host species considered

Free-ranging snakes were captured from March 2020 to June 2022 across 10 countries: Portugal, Spain, France, Switzerland, Germany, Austria, Czech Republic, Hungary, Poland, and Ukraine. The number of sites where snakes were collected ranged from one to nine per country, for a total of 43 sites (Fig. 1, Table S1). Sites were selected based on preexisting geopolitical boundaries within each country (e.g., counties or regions), to account for potential large variation in landscape and geography within each country.

**Figure 1.**
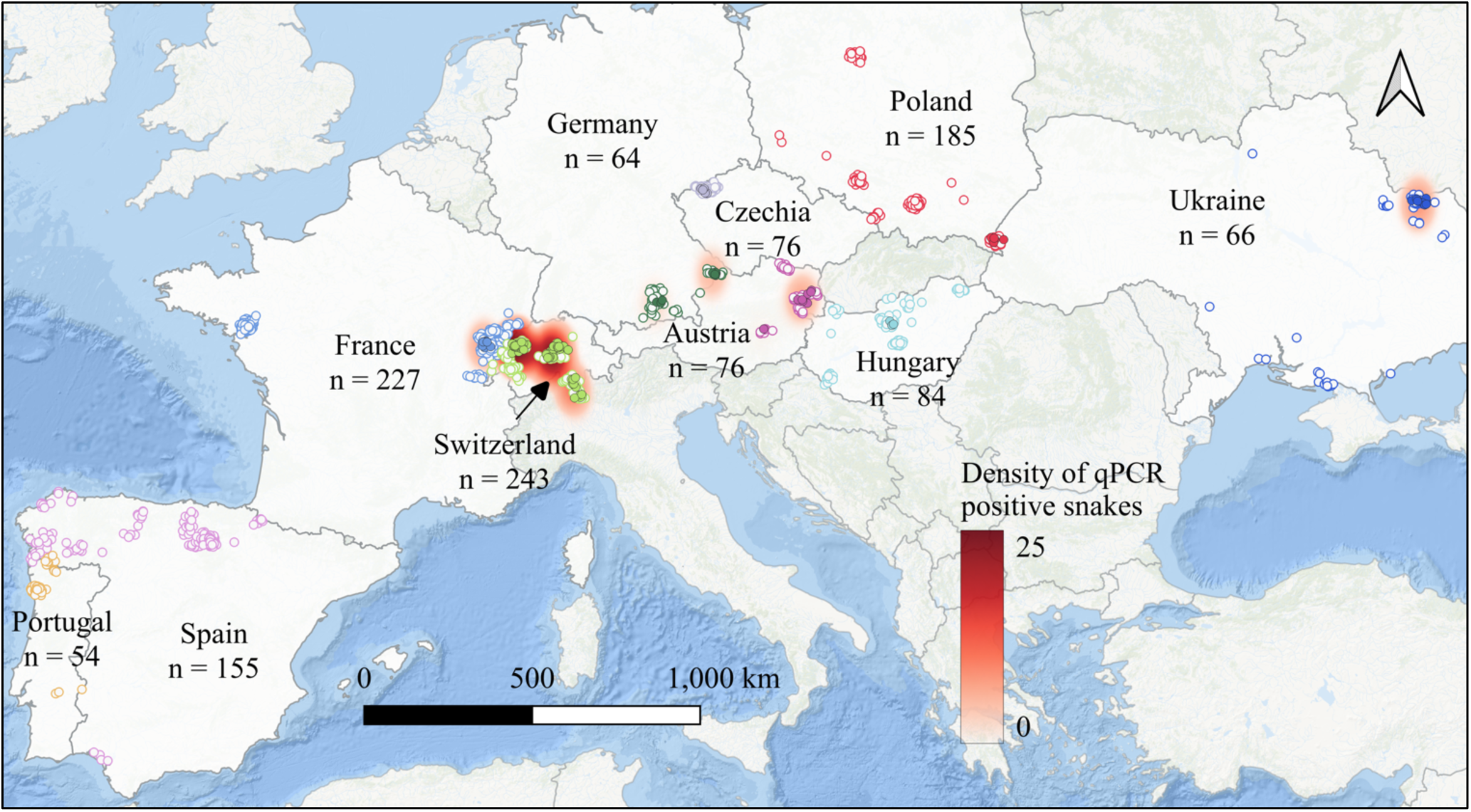
Spatial distribution of snake captures and detections of *O. ophidiicola* across Europe. Each circle represents an individual snake capture and overlapping points were slightly jittered for visualization. Different colors are used to distinguish countries, filled points indicate snakes that were qPCR positive, and outlined points are qPCR negative snakes. Underlying density heatmap shows spatial distribution of *O. ophidiicola* infection risk based on qPCR positive detections using a kernel density estimation algorithm for visualization. We used a 100-km radius around each positive point and the scale bar indicates point density (i.e. relative disease risk) across each region.

### (b) Capture and sampling

Snakes were located by visual encounter surveys, an approach frequently used for sampling snakes (Dorcas and Willson 2009). Surveys were guided by pre-existing knowledge of snake presence or prediction of suitable habitats to sample as many species and individuals as possible, and in a wide variety of habitat types. Snakes were captured by hand, placed in individual cloth bags for temporary holding during processing and sampling, and released at their capture location. Sterile handling procedures, including frequent glove changes and decontamination of gear between each snake were followed during sample collection to avoid cross-contamination. Snakes were individually identified using photo-identification or marking (using PIT tags or scale-clipping). For each snake captured, location and morphometric data were collected including latitude/longitude, species, sex, snout-vent-length, tail length, and weight.

Snakes were swabbed in duplicate (except for a few individuals that were swabbed only once due to limitations in the field) using a pre-moistened, sterile polyester-tipped applicator (Puritan^®^, Guilford, Maine, USA) by running the swab five times (back and forth counting as a single pass) on the ventral and dorsal areas (from the neck down to the vent), and two times on the face of the snake. If a skin lesion was observed, a separate swab was used to specifically swab the lesion and skin immediately adjacent to it by rubbing the swab over the affected skin. In addition, for all snakes that had visible lesions, we collected photos for later quantification of infection severity (see below). Swab tips were individually stored in a 2 mL sterile tube in a cooler with ice while in the field, and later stored frozen at -20°C until analysis.

### (c) Sample extraction and qPCR

A total of 2628 swabs were collected and processed by one of two laboratories following the exact same methods as described below. DNA was extracted from swabs using 250 µL of PrepMan® Ultra Sample Preparation Reagent (Life Technologies, Carlsbad, California, USA) with 100 mg of zirconium/silica beads, following a previously published protocol (Hyatt et al. 2007). Briefly, samples were homogenized for 45 sec in a bead beating grinder and lysis system (MP Biomedicals, Irvine, California, USA) and centrifuged for 30 sec at 13000 g to settle all material to the bottom of the tube. Homogenization and centrifugation steps were repeated, and tubes were incubated at 100°C in a heat block for 10 min. Tubes were then cooled at room temperature for 2 min, then centrifuged for 3 min at 13000 g. Fifty to 100 µL of supernatant was recovered and stored at -80°C. Extraction blanks (negative controls) were prepared using 250 µL of PrepMan® Ultra Sample Preparation Reagent and 10 mg of zirconium/silica beads only.

Quantitative PCR targeting the internal transcribed spacer region (ITS) specific to *O. ophidiicola* was performed on a real-time PCR QuantStudio 5 (Thermofisher Scientific, Waltham, Massachusetts, USA) (Bohuski et al. 2015). QuantiFast Master Mix (QuantiFast Probe PCR + ROX vial kit, Qiagen, Germantown, USA) was prepared according to manufacturer’s recommendations for a final reaction volume of 25 µL, which included 5 µL of extracted DNA. Cycling conditions were as follows: 95°C for 3 min, then 95°C for 3 sec and 60°C for 30 sec for a total of 40 cycles. For each plate run, a negative control (water added instead of extracted DNA) and a 6-point (each point run in triplicate) standard curve using synthetic double-stranded DNA (gBlock, Integrated DNA Technologies, Coralville, Iowa) of the target region (1.0 x 10^2^, 1.0 x 10^1^, 1.0 x 10^0^, 1.0 x 10^-1^, 1.0 x 10^-2^, 1.0 x 10^-3^ fg/µL) were included. Samples that were positive were analyzed in duplicate, and a snake was determined to be positive if any swab associated with that snake was positive by qPCR.

### (d) Sequencing and genotyping

Samples in which *O. ophidiicola* was detected by qPCR were subjected to follow up genotyping analysis. We targeted a portion of the internal transcribed spacer 2 (ITS2) for this analysis because the ITS2 exhibits variability between previously described clades of *O. ophidiicola* and because ITS2 is a multicopy gene that can be amplified from samples containing very small amounts of *O. ophidiicola* DNA (it is also the target of the qPCR assay). We used a nested PCR protocol that consisted of first amplifying the entire ITS2 region with the panfungal primer ITS3 and ITS4 (White et al. 1990). The first reaction consisted of 10 µL of 2x QuantiNova probe PCR master mix (Qiagen, Venlo, Netherlands), 3.9 µL of molecular grade water, 0.5 µL of each primer (20 µM each), 0.1 µL of 20µg/µL bovine serum albumin, and 5 µL of DNA extracted with the PrepMan procedure described above. Cycling conditions were as follows: 95°C for 3 min; 40 cycles of 95°C for 10 sec, 56°C for 30 sec, and 72°C for 30 sec; final extension at 72°C for 5 min. For the second reaction, primers ITS3 and Oo-rt-ITS-R (Bohuski et al. 2015) were used. Each reaction consisted of 0.5 µL of the PCR product from the first reaction added to 13.375 µL molecular grade water, 5 µL of GoTaq Flexi buffer (Promega Corporation, Madison, Wisconsin, USA), 2 µL of dNTPs (2.5 mM each), 1.5 µL of 25 mM MgCl_2_, 1.25 µL of each primer (20 µM each), and 0.25 µL of GoTaq polymerase. Cycling conditions for the second PCR were: 95°C for 10 min; 45 cycles of 95°C for 30 sec, 56°C for 30 sec, and 72°C for 1 min; final extension at 72°C for 5 min. Products from the second PCR were visualized on an agarose gel, and those containing bands were sequenced in both directions using the Sanger method with primers ITS3 and Oo-rt-ITS-R.

Samples that generated messy chromatograms or appeared to contain single nucleotide polymorphisms (SNPs) indicative of multiple *O. ophidiicola* genotypes were re-amplified with the second PCR using a proofreading polymerase (15.75 µL of molecular grade water, 5 µL of 5x SuperFi buffer [Thermo Fisher Scientific Corporation, Waltham, Massachusetts, USA], 2 µL of dNTPs (2.5 mM each), 0.625 µL of 20 µM each primer, 0.5 µL of Platinum SuperFi DNA polymerase [2U/µL], and 0.5 µL of product from the first PCR; cycling conditions were the same as described for the second reaction above). The resulting amplicons were cloned using the Invitrogen Zero Blunt TOPO PCR cloning kit for sequencing (Thermo Fisher Scientific Corporation, Waltham, Massachusetts, USA), and individual transformants were sequenced.

Individual ITS2 sequences generated in our study were assigned to genotypes. Sequences with 100% identity across the ITS2 region of *O. ophidiicola* were considered to be the same genotype; any sequence differing from another by at least one SNP was classified as a unique genotype.

### (e) Quantification of disease severity

Disease severity was measured by calculating the percentage of surface area of each snake covered by lesions. Using the image processing program ImageJ (Schneider et al. 2012) and the photos of the snakes taken in the field, we measured each lesion five times and recorded the mean length and width. We calculated the surface area of each lesion on a particular snake and added up the surface area of each lesion to determine the total lesion surface area. Using the morphometric measurements collected in the field, we also calculated each snake’s total surface area from the snout to the tail tip. We then calculated the percentage of total surface area covered by lesions. We also quantified disease severity following a previously described SFD scoring system (Baker et al. 2019) using a combination of lesion type, location, number, and coverage.

Scores ranged from 4 (mild) to 12 (most severe). We compared the relationship between lesion score and percentage of total surface area covered by lesions (Fig. S1) and used lesion surface area as a more quantitative measure of severity as opposed to the ordinal metric in the final analyses to describe disease severity.

### (f) Statistical analyses

We analyzed data using Bayesian hierarchical models and we assessed statistical support using credible intervals that do not overlap zero. We fit all models (unless otherwise noted) using the No-U-Turn Sampler (NUTS), an extension of Hamiltonian Markov chain Monte Carlo (HMCMC). We created all Bayesian models in the Stan computational framework (http://mc-stan.org/) accessed with the “brms” package (Bürkner 2017). To improve convergence and avoid over-fitting, we specified weakly informative priors (a normal distribution with mean of zero and standard deviation of 10), unless otherwise noted. Models were run with a total of 4 chains for 2000 iterations each, with a burn-in period of 1000 iterations per chain resulting in 4000 posterior samples, which, given the more efficient NUTS sampler, was sufficient to achieve adequate mixing and convergence. All 𝑅̂ values were less than or equal to 1.01, indicating model convergence. We performed all statistical analyses in R software version 4.2.0 (R Core Team 2022).

To examine hotspots (i.e., relative disease risk) of the pathogen *O. ophidiicola* across Europe, we used the Getis-Ord Gi* analysis with Local Indicators of Spatial Association (LISA) statistics in the open-source Geographic Information System QGIS (version 3.22.10). A grid map of 100 km x 100 km was created across the landscape to aggregate point data and look at significant spatial clustering of neighboring features (each square) using the resultant z-scores and p-values. A heatmap was created to visualize the *O. ophidiicola* infection risk zone using a kernel density estimation with a quartic kernel shape and kernel radius of 100 km in QGIS. To investigate the effect of host species on pathogen prevalence, we used a Bayesian multilevel model with a Bernoulli distribution, pathogen detection as our response variable (0|1), species as our predictor variables, and a group-level effect of site. We excluded any species that had been sampled fewer than four times. We ran a similar model as described above with genus instead of species and included species as a group-level effect.

To investigate differences in lesion prevalence and disease severity in snakes that were qPCR positive, we first ran a multilevel model with a Bernoulli distribution, with detection of lesions on a snake in the field (0|1) as our response variable, species as our predictor variable, and a group-level effect of site. To examine differences in disease severity for the snakes that were positive for *O. ophidiicola,* our response variable was each square millimeter of surface area of the snake, which was treated as Bernoulli trials in a binomial sample (1 = lesion present or 0 = no lesion), and our group-level effects were individual snake and site. We performed a comparison between our lesions severity metric (percent of body covered in lesion) and previously described lesion scoring system (Baker et al. 2019), using a hierarchical model with the scoring system as our response variable, and the fraction of skin covered in lesions as our predictor with a Poisson distribution.

We examined the best model that explained *O. ophidiicola* prevalence across the landscape using leave-one-out cross-validation (LOO). The multilevel models included the population level effects of just species, just pathogen clade, an additive model of species and clade, and an interactive model of species and clade, with a Bernoulli distribution, pathogen detection as our response variable (0|1), and with site as a group-level effect for all four models. Models were run with a total of 4 chains for 6000 iterations each, with a burn-in period of 1500 iterations per chain resulting in 18000 posterior samples. We pooled the four genotypes together (i.e., I-A, I-B, II-D/E and II-F) into two clades (clade I and II) as sample sizes were generally too small across species to look at separately. The clade variable consisted of either clade I (sites where only snakes infected with clade I were detected) or both clades I and II (sites where snakes were infected with either clade I or clade II, as there were only a few locations with detections of only clade II). The final clade dataset used for this analysis included 16 sites and 751 snakes.

Prevalence by pathogen clade comparisons are only reported for four species (*N. natrix*, *N. tessellata*, *N. helvetica*, and *Z. longissimus*) for which there was sufficient data. To further examine the contribution of host species and pathogen clade to disease hotspots we also ran the analysis as described above with the addition of a spatial conditional autoregressive (CAR) term in “brms”. Finally, to examine how pathogen clade influenced infection severity, we performed an analysis where our response variable was each square millimeter of surface area of the snake, which was treated as a Bernoulli trial in a binomial sample and our predictor was pathogen clade, with a group-level effect of individual snake and species.

## 6. Results

We captured 1254 individual snakes from 21 species representing 6 genera (Fig. 1, Table S1). A total of 2628 swabs were collected, including 2357 full body skin swabs and 271 lesion swabs. Overall, *O. ophidiicola* prevalence confirmed by qPCR was 8.7% (n = 109 positive snakes) and prevalence was highly variable across the landscape. The hotspot analysis confirmed that sites in Switzerland and across the border in France (Franche-Comte) have elevated pathogen prevalence compared to surrounding areas (99% confidence interval, Z-score ≥ 2.58; p ≤ 0.01, Fig. 1 and S2), as well as sites sampled in southern Germany, eastern Austria, and eastern Ukraine, but to a lesser extent (95% confidence interval, 1.96 ≤ Z score < 2.58; 0.01 < p ≤ 0.05). Other locations where *O. ophidiicola* was detected (Czech Republic, Hungary, and Poland) were not found to be significant hotspot (p > 0.05). Pathogen prevalence was highest in Switzerland (26.7%, Fig. 1, S3), followed by Germany (12.5%, Fig. S3) and Ukraine (12.1%, Fig. S3). Poland and the Iberian Peninsula (Spain and Portugal) had the lowest prevalence at 2.7%, 0.0%, and 0.0% respectively, despite comparable sample sizes to other locations (Table S1).

In addition to geographic variation, differences in *O. ophidiicola* prevalence were considerable among hosts (Fig. S4). We found statistical support (results formatted as coefficient ± standard deviation (95% credible intervals)) that species in the *Natrix* genus had higher prevalence (8.2%; intercept: -2.51 ± 0.50 (−3.62, -1.67)) than other genera sampled (0.7%, *Coronella* coeff: -3.13 ± 1.23 (−6.07, -1.24); 0.3%, *Dolichophis* coeff: -8.64 ± 5.60 (−22.53, - 0.99); 0.2%, *Vipera* coeff: -4.05 ± 0.87 (−5.95, -2.53)), except for *Hierophis* and *Zamenis* (4.2% and 5.9 %, coeffs: -0.81 ± 0.48 (−1.79, 0.10) and -0.41 ± 0.43 (−1.27, 0.39), respectively). There was also large variation in pathogen prevalence among congeneric species. *Natrix tessellata* had higher *O. ophidiicola* prevalence (model prediction: 15.8 ± 6.2%; Fig. 2 and Table S2) compared to several other members of the *Natrix* genus including *Natrix astreptophora* (0.6%, coeff: -8.5 ± 5.8 (−21.77, -0.39)), and *Natrix maura* (2.0%, coeff: -2.66 ± 1.31 (−5.7, -0.56), but there was no statistical support for differences from *Natrix helvetica* (13.7%, coeff: -0.16 ± 0.38 (−0.9, 0.58)) or *Natrix natrix* (7.6%, coeff: -0.75 ± 0.49 (−1.73, 0.23)); Fig. 2 and Table S2).

**Figure 2.**
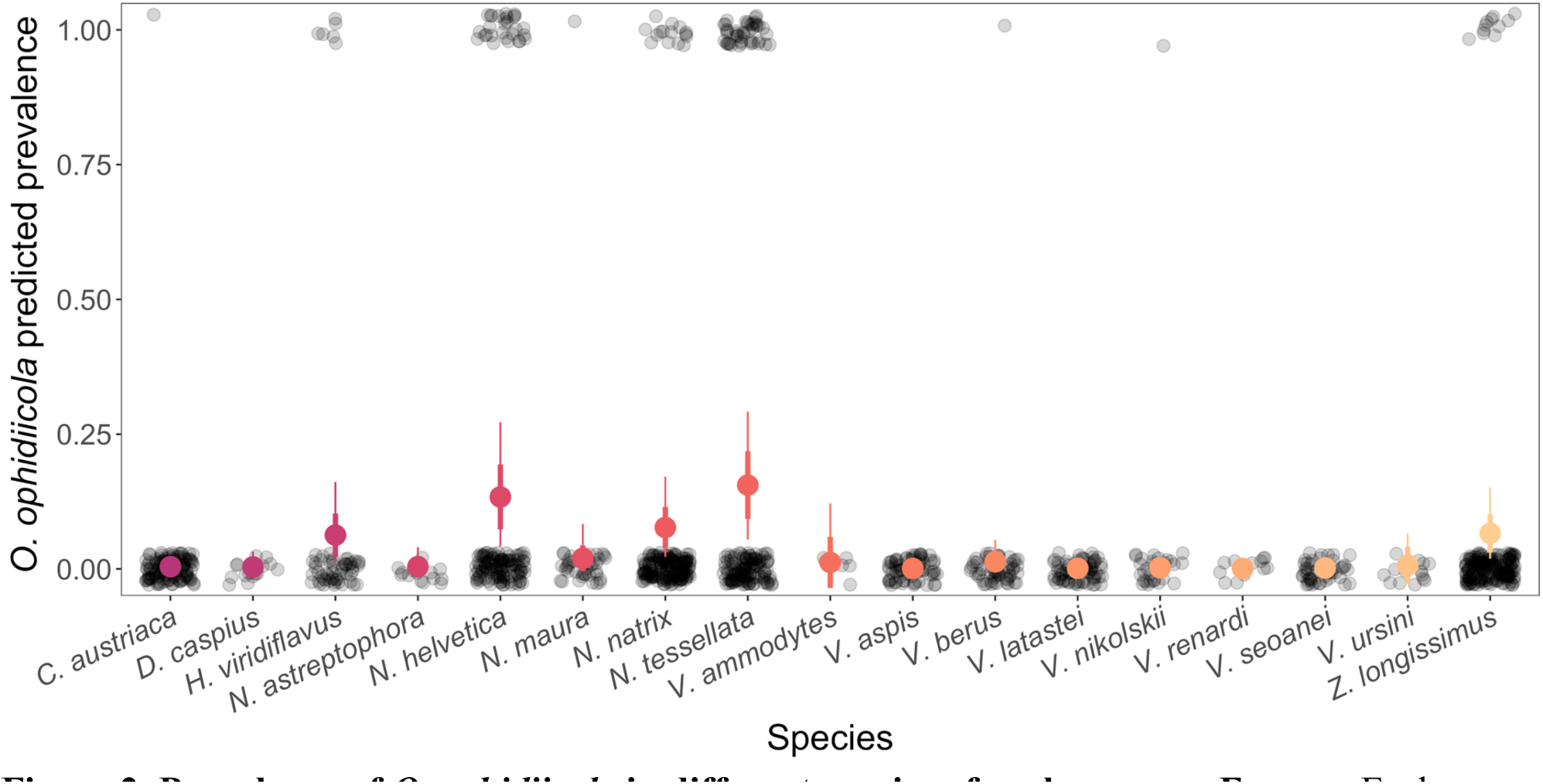
Prevalence of *O. ophidiicola* in different species of snakes across Europe. Each black circle represents a single snake as being either negative (0) or positive (1), which was then used to calculate pathogen prevalence (fraction of the population that was positive). Larger circles and whiskers show the model predicted posterior mean ± standard deviation (thick lines), and 95% credible intervals (thin lines) for each species across all countries. Colors indicate different species.

Overall, skin lesions were observed in 187 snakes from 15 species across all countries, but only 46.5% of those tested positive by qPCR (n = 87). Of all the snakes that tested positive by qPCR (n = 109), 80% of those had skin lesions that were consistent with SFD, while the other 20% had no visible skin lesion (n = 22) (Fig. 3a, Fig. S5). We found a high probability of finding lesions on a snake if they tested positive for *O. ophidiicola* (range 46.2% – 92.8%, except for two viper species which had no visual sign of disease, Fig. 3a). We observed variation in disease severity but no statistical support for differences among species (Fig. 3b, Table S3).

**Figure 3.**
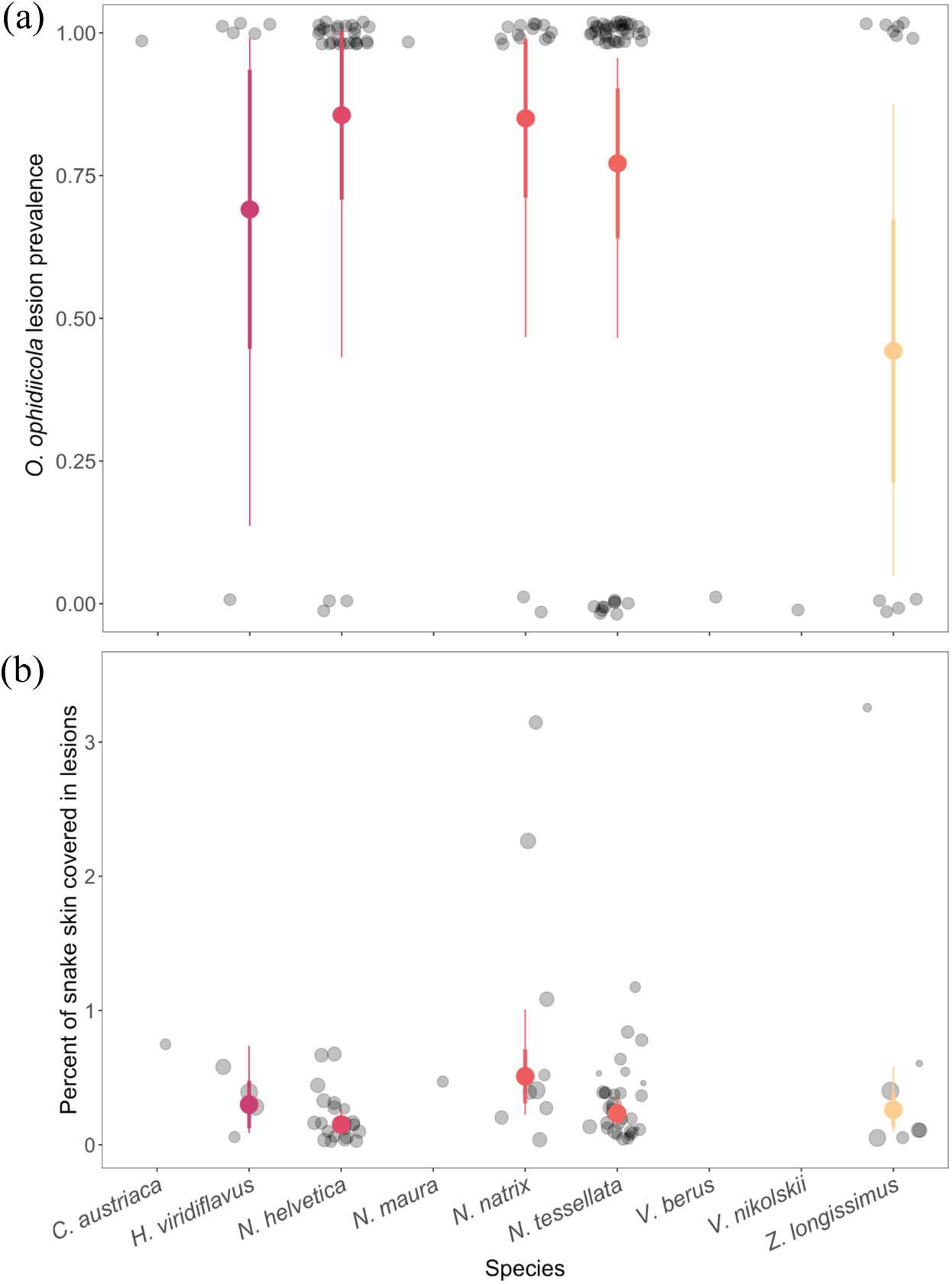
Lesion prevalence (a) and disease severity (b) in *O. ophidiicola*-positive snakes across different species. Color circles and whiskers show the model predicted posterior mean, ± standard deviation (thick lines), and 95% credible intervals (thin lines) for different species across all countries. (a) Each black circle represents a single snake as being either negative (0) or positive (1) for presence of lesions, which was used to calculate the proportion of the population that tested positive (prevalence). (b) Each black circle represents the percentage of the body of a single snake covered in lesions and the size of the circle is proportional to the total surface area of the snake (scale ranges 250 – 1,000 cm^2^).

Genotyping analyses were successful for 85.3% of positives swabs (93 total swab samples) that were qPCR positive for *O. ophidiicola*. A total of four unique genotypes were observed, belonging to two of the major *O. ophidiicola* clades (clade I & II, (Ladner et al. 2022)) (Table S4). Two of these genotypes (designated here as I-A and I-B) resided within clade I (i.e., the “European clade”). What we refer to as genotype I-A had an ITS2 sequence identical to strains previously isolated from Great Britain, while genotype I-B had an ITS2 region sequence identical to a strain from Czechia (Franklinos et al. 2017). The remaining two genotypes that we observed in our study resided within clade II (i.e., the “North American clade”) and were identical to ITS2 region sequences of clonal lineages II-D/E (lineages D and E have identical sequences in the ITS2 region) and II-F (Ladner et al. 2022). Here we refer to these genotypes as II-D/E and II-F, respectively, although strains detected in this study may not be true representatives of the clonal lineages reported from North America since recombinant strains can have identical ITS2 sequences as clonal lineages (Ladner et al. 2022). We found that both clade I and clade II are present across much of continental Europe, ranging from eastern France to eastern Ukraine; however, both clades were not present in all locations where *O. ophidiicola* was detected. Genotype I-A was detected primarily in western Europe (Switzerland, Germany, and Austria), whereas genotype I-B was detected in eastern Europe (Czechia, Austria, Hungary, Poland, and Ukraine) (Fig. 4a). Genotype II-D/E was more widely distributed across Europe, whereas genotype II-F was only found along a single lake in Switzerland (Fig. 4a). On two occasions, snakes were found to be infected with multiple genotypes of *O. ophidiicola*: a snake from Switzerland from which genotypes I-A and II-D/E were detected and a snake from Hungary from which genotypes I-A and I-B were detected.

**Figure 4.**
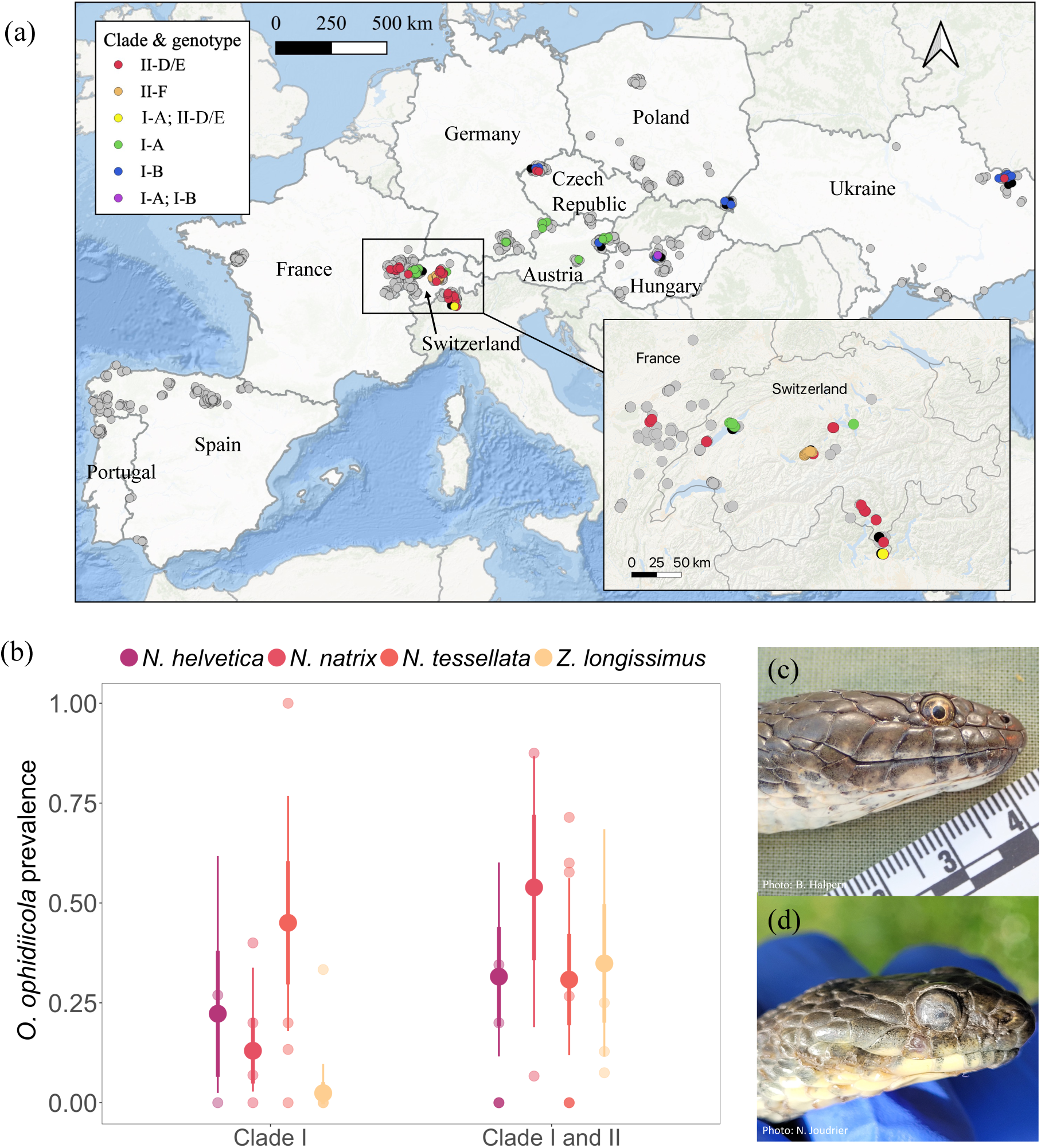
Distribution of *O. ophidiicola* clades across Europe. (a) *O. ophidiicola* clades and lineages from positive detections across the landscape in Europe. Points are slightly jittered for better visual representation of the sampling range. Color of the point indicates the clades and genotypes, samples that were qPCR negative are represented as grey points, and samples that failed to amplify with the genotyping PCR are represented as black points. Yellow and purple points represent simultaneous detections of genotypes I-A and II-D/E and genotypes I-A and I-B, respectively, from the same swab sample (i.e., snakes infected with multiple genotypes). The enlarged map (inset) shows better resolution of detections in Switzerland. (b) *O. ophidiicola* prevalence across snake species based on *O. ophidiicola* clade presence at the different sites.

Small color points are mean prevalence at a site for a given species and clade. Large color points and whiskers show the model predicted posterior mean, ± standard deviation (thick lines), and 95% credible intervals (thin lines) across different species. (c) Photo of a *N. tessellata* from Hungary infected with *O. ophidiicola* from clade I-B. (d) Photo of a *N. tessellata* from Switzerland infected with *O. ophidiicola* from clade II-F showing facial infection.

The top two models, as determined through LOO, that best predicted *O. ophidiicola* prevalence across the landscape included both host species and pathogen clade as predictor variables, with the best fit model including an interaction between these two variables (Table S5). There was statistical support that *N. natrix* and *Z. longissimus* have higher probabilities of being infected when clade II was detected at a site compared to just clade I (*N. natrix* coeff: 2.09 ± 1.09 (0.05, 4.27), *Z. longissimus* coeff: 3.15 ± 1.23 (0.84, 5.69)). A third species, *N. helvetica*, was associated with higher probability of infection when clade II was detected at a site, but the credible intervals included zero (coeff: 0.64 ± 1.16 (−1.59, 3.03)). Conversely, we found that the species with the highest pathogen prevalence, *N. tessellata,* had no differences in infection probability based on clade (Fig. 4b, Table S6). Results from the spatial regression resulted in qualitatively similar results, where both species present and pathogen clade were important predictors of pathogen prevalence (*N. natrix* coeff: 1.10 ± 0.75 (−0.73, 2.35), *Z. longissimus* coeff: 1.74 ± 0.85 (0.10, 3.44), *N. helvetica* coeff: 0.09 ± 0.44 (−0.72, 0.95)), except for *N. tessellata* (coeff: -0.59 ± 0.54 (−1.66, 0.42)) (Table S7). Snakes that were infected by a strain of *O. ophidiicola* belonging to clade I generally had less severe disease when compared to snakes infected with a strain from clade II, although there was no strong statistical support as the credible intervals overlapped zero (clade I coeff: -0.10 ± 0.34 (−0.77, 0.57), Fig. S6).

## 4. Discussion

Our results support the presence of hotspots for SFD across Europe and the potential factors that contribute to higher infection prevalence. *O. ophidiicola* was detected in all countries studied except for those on the Iberian Peninsula (Spain and Portugal), which may be attributed to numerous factors (e.g. different species community, geographical barriers (Pyrenees mountains), or environmental conditions) that may be unsuitable for persistence and growth of *O. ophidiicola*. Our surveys establish four regions with elevated prevalence of *O. ophidiicola* in Europe, with the highest being in Switzerland.

Snake fungal disease has garnered much attention over the last few decades, as this disease has negatively affected snake populations (Allender et al. 2015, Lorch et al. 2016). Despite this, few studies have systematically examined differences in disease severity and prevalence across species. We found high variability of infection prevalence among snake species, irrespective of the sampling location. Snakes in the *Natrix* genus had a higher probability of infection compared to other genera, indicating that species in this genus may be more susceptible to *O. ophidiicola* infections and are likely to be important in maintaining elevated pathogen prevalence in a region. Underlying host characteristics such as dependence on aquatic habitats, have previously been found to be associated with higher *O. ophidiicola* infection prevalence (McKenzie et al. 2019), which could partly explain increased infection in this genus. *Natrix tessellata* had the highest pathogen prevalence, followed by *N. helvetica* and *N. natrix*, with all three species being either semiaquatic or living near water (with *N. tessellata* being more piscivorous). Interestingly, only two vipers (out of a total of 341 samples) tested positive for *O. ophidiicola*, and both snakes had no visual signs of infection (i.e. no lesions present). This indicates that viperids may not be competent hosts for *O. ophidiicola*, possibly due to environmental associations or behavioral and physiological mechanisms. Contrary to this, North American pit vipers, such as the massassauga rattlesnake (*Sistrurus catenatus*), have been reported to develop severe clinical signs of SFD (Allender et al. 2011). This could be attributed to a sampling bias from more intensive monitoring of threatened rattlesnake species, or rattlesnakes may have increased susceptibility to SFD due to their habitat requirements (massassauga rattlesnakes are found near swamps and marshes) compared to European vipers, which are generally associated with drier environments (except for *V. berus* which can be associated with cool and humid habitats).

In our study, no mortality was reported, and snakes generally appeared healthy except in a few cases where infection was severe and had spread to the face with possible disruption to foraging behavior (Fig. S4). The low disease severity observed in Europe could be the result of increased host resistance or generally lower pathogen virulence. We also found that only 46% of snakes with lesions tested positive for *O. ophidiicola*, which has also been reported from North America (Chandler et al. 2019, Haynes et al. 2020). The lesions that could not be attributed to *O. ophidiicola* infection looked similar to SFD skin lesions and may be fungal or bacterial in origin (Dubey et al. 2022). Further research investigating other sublethal effects of SFD and the interaction between *O. ophidiicola* and other pathogens is important.

We found that models accounting for both host species and pathogen clade best explained the variation in pathogen prevalence across the landscape. Importantly, the top model included an interaction between host species and pathogen clade, indicating that the effect of clade is not the same across species. Clade II was found in two of the four pathogen hotspots across Europe, and generally when clade II was present in an area, we found support that the probability of detecting *O. ophidiicola* was higher for three (*N. natrix*, *Z. longissimus*, and *N. helvetica)* of the four species with the highest prevalence. Hotspots in Switzerland were primarily driven by *N. tessellata*, which showed no difference in detection based on whether just clade I or both clades I and II were present. This indicates that *N. tessellata* may be equivalently susceptible to both clades I and II or that other factors may be contributing to infection probability in this species.

The history and origin of *O. ophidiicola* in Europe is unclear. Sampling of museum specimens has revealed that strains of *O. ophidiicola* belonging to clades I and II were present in Switzerland as early as 1959 (Origgi et al. 2022). It is estimated that clade I shared a common ancestor within the last 100 to 500 years, but that analysis only included four clade I strains and may greatly underestimate the time that *O. ophidiicola* has been present in Europe (Ladner et al. 2022). Despite the designation of clade II as the “North American” clade, it is believed that *O. ophidiicola* is not native to North America and multiple introductions (most likely from Eurasia) have occurred over the last century (Ladner et al. 2022). Thus, it is plausible that either or both clades I and II are native to Europe. Strains of *O. ophidiicola* isolated from wild snakes in Taiwan reside within clade II (Sun et al. 2021, Ladner et al. 2022), which could also indicate a southeast Asian origin for that clade, raising the possibility that clade II is not native to Europe.

Clade II has been detected on captive snakes in Europe (Ladner et al. 2022), which could serve as a source for transmission into wild populations. An introduction of clade II into Europe sometime before 1960 and subsequent spread could explain the wide distribution of this clade as detected in our sampling. We detected two genotypes within clade II. One of these (II-D/E) was widely distributed, whereas the other (II-F) was detected in a single snake community around a lake in Switzerland. That snake community includes an introduced population of *N. tessellata*. Taken together, this could indicate that genotype II-F was more recently introduced to Europe, perhaps through the release of snakes originating in captivity. However, determining the genetic diversity and origin of the various lineages of *O. ophidiicola* in Europe would require more in-depth studies.

We find several disease hotspots in Europe, which could be attributed, at least partially, to specific host species and the presence of distinct pathogen clades. The shape of this relationship varied among areas with higher disease prevalence, and in some cases, host species had higher infections regardless of pathogen genotypes, and others were likely attributed to the presence of specific susceptible hosts infected with a pathogen clade that may be more transmissible. Although virulence is recognized as an important factor in the effects of disease on host populations, the general lack of landscape level data on pathogen lineage distribution and association with disease has likely limited our ability to determine its importance for other disease hotspots.

## Ethics

Handling of snakes was reviewed and approved by Virginia Tech Institute for Animal Care and Use Committee protocol 20-055. Permits to conduct our field study were obtained when necessary, and were granted by the Regional office of the Karlovarian region, Department of Environment and Agriculture in Czech Republic (permit # KK/1098/ZZ/20-4), the General and Regional Directorates of Nature Conservation in Poland (permit # DZP-WG.6401.91.2020.TŁ; DZP-WG.6401.91.2020.TŁ.2, WPN.6401.270.2019.MF, WPN.6401.17.2020.KW.2, WPN.6401.9.2021.KW.2), Portuguese and Spanish wildlife legislation (permits # 295/2020/CAPT, 146/2021/CAPT, 201999902471003/IRM/MDCG/mes, EB-018/2020, EB-015/2021, AUES/CYL/192/2020, AUES/CYL/54/2021, A/2021/036, 0001-0261-2021-000003), the Direction régionale de l’environnement of Franche-Comte (ONAGRE 2020-01-17-00122; 25-2022-02-17-00001) and Loire-Atlantique (Cerfa 13616, permit # 64/2016) in France, the Museum of Natural History in Budapest, Hungary (permit # PE-KTFO/1568-18/2020), Austrian legislations (permits # MA22 – 1089768-2020, RU5-BE-64/023-2022, RU5-BE-64//022-2021, ABT13-53W-50/2018-2), the Federal Office for the Environment (FOEN) in Switzerland with veterinary authorization (VD3718, ID 33612), and cantonal authorizations (Neuchatel: FS-08/2021, Grisons: AV-2022-338, Vaud: 2021-3537).

## Acknowledgements

We thank Otto Aßmann, Florian Bacher, Jacek Błachuta, Philomin Briot, Jon Buldain, Maxime Chèvre, Pierre Cheveau, Sylvain Dubey, Anna Egerer, Lea Endrejat, Karin Ernst, Niklas Franzen, Inês Freitas, Georg Gassner, Davy Guinchard, Krisztián Harmos, Marietta Hengl, Piotr Hubal, Kacper Jurczyk, Yurii Kornilev, Roman Kurek, Nahla Lucchini, Adrian Neumann, Daniel Renner, Sandra Wallner, and Bartłomiej Zając for assistance in the field. Many thanks also to Kate Langwig for statistical insights, and to Oleksandra Klynova and Megan Winzeler for laboratory assistance.

## Author contributions

GB: conceptualization, data curation, investigation, formal analysis, visualization, methodology, writing—original draft, writing—review and editing; JML: methodology, investigation, writing—review and editing; NJ: methodology, resources, investigation, writing—review and editing; SB: investigation, resources, writing—review and editing; TC: methodology, investigation, writing—review and editing; MF: investigation, writing—review and editing; FM-F: investigation, writing—review and editing; GG: investigation, writing—review and editing; BH: investigation, writing—review and editing; AK: investigation, writing—review and editing; KK: investigation, writing—review and editing; OL: investigation, writing—review and editing; AM: methodology, investigation, writing—review and editing; RM: investigation, writing— review and editing; SS: investigation, review and editing; BS: investigation, writing—review and editing; SU: methodology, resources, investigation, writing—review and editing; OZ: investigation, writing—review and editing; JRH: conceptualization, resources, supervision, funding acquisition, writing—review and editing.

## Disclaimer

Any use of trade, firm, or product names is for descriptive purposes only and does not imply endorsement by the U.S. Government.

**<u>Data availability statement:</u>** The datasets and code generated for this study will be made available in Dryad Digital Repository upon final submission.

