## Supplemental material for "Contribution of host species and pathogen clade to snake fungal disease hotspots in Europe"

**Supporting information**

**Table S1.** Species and sample sizes across all 10 countries sampled from March 2020 to June 2022 for the presence of *Ophidiomyces ophidiicola*, the pathogen responsible for Snake Fungal Disease. The 4-letter codes represent genus and species for each snake species sampled as follows: *Coronella austriaca* (COAU), *Coronella girondica* (COGI), *Dolichophis caspius* (DOCA), *Elaphe sauromates* (ELSA), *Hierophis viridiflavus* (HIVI), *Malpolon monspessulanus* (MAMO), *Natrix astreptophora* (NAAS), *Natrix helvetica* (NAHE), *Natrix natrix* (NANA), *Natrix maura* (NAMA), *Natrix tessellata* (NATE), *Vipera ammodytes* (VIAM), *Vipera aspis* (VIAS), *Vipera berus* (VIBE), *Vipera latastei* (VILA), *Vipera nikolskii* (VINI), *Vipera renardi* (VIRE), *Vipera seoanei* (VISE), *Vipera ursini* (VIUR), *Zamenis longissimus* (ZALO), *Zamenis scalaris* (ZASC). For each species, the numbers within each cell represent: total number of snakes with skin lesion / total number of snakes testing positive by qPCR / total number of snakes sampled. The size of each site is reported in km^2^.

| ***AUSTRIA (n = 76)*** |  | *Species* | | | | |  |  |  |  |
| --- | --- | --- | --- | --- | --- | --- | --- | --- | --- | --- |
| Site | Size | COAU | NANA | NATE | VIAM | ZALO |  |  |  |  |
| Lower Austria | 19186 | 1/0/3 | 3/0/26 | 0/4/20 | 0/0/7 | 1/0/8 |  |  |  |  |
| Styria | 16401 | - | 0/1/5 | - | - | 0/0/4 |  |  |  |  |
| Vienna | 415 | - | - | 2/3/3 | - | - |  |  |  |  |
| ***CZECH REPUBLIC (n = 100)*** |  | *Species* | | | | |  |  |  |  |
| Site | Size | COAU | NANA | NATE | VIBE | ZALO |  |  |  |  |
| Karlovy Vary | 3315 | 4/1/19 | 2/1/15 | 0/0/17 | 0/0/9 | 3/3/40 |  |  |  |  |
| ***FRANCE (n = 227)*** |  | *Species* | | | | | | |  |  |
| Site | Size | COAU | HIVI | NAHE | NAMA | VIAS | VIBE | ZALO |  |  |
| France-Comte | 16202 | 5/0/14 | 3/2/18 | 1/0/14 | 1/0/25 | 0/0/16 | 0/0/10 | 10/5/39 |  |  |
| Loire-Atlantique | 6881 | 0/0/2 | - | 0/0/25 | - | 0/0/10 | 0/0/14 | 0/0/40 |  |  |
| ***GERMANY (n = 64)*** |  | *Species* | | | | | |  |  |  |
| Site | Size | COAU | NAHE | NANA | NATE | VIBE | ZALO |  |  |  |
| Upper/Lower Bavaria | 27859 | 0/0/3 | 2/0/3 | 1/2/29 | 4/5/11 | 1/1/15 | 2/0/3 |  |  |  |
| ***HUNGARY (n = 84)*** |  | *Species* | | | | | | |  |  |
| Site | Size | COAU | DOCA | NANA | NATE | VIBE | VIUR | ZALO |  |  |
| Bacs-Kiskun | 8445 | 1/0/3 | - | 0/0/5 | - | - | 0/0/8 | - |  |  |
| Borsod-Abauj-Zemplen | 7250 | - | - | 0/0/2 | 0/0/5 | - | - | - |  |  |
| Budapest | 525 | 0/0/1 | 1/0/12 | - | 4/2/15 | - | - | 0/1/3 |  |  |
| Gyor-Moson-Sopron | 4208 | - | - | - | - | - | 0/0/6 | - |  |  |
| Heves | 3637 | - | - | 0/0/2 | - | - | - | - |  |  |
| Nograd | 2545 | 0/0/3 | - | 0/0/1 | - | - | - | 0/0/1 |  |  |
| Pest | 6393 | - | - | 0/0/2 | 0/0/4 | - | - | - |  |  |
| Somogy | 6065 | - | - | 0/0/5 | - | 0/0/5 | - | - |  |  |
| Vas | 3336 | - | - | - | - | 0/0/1 | - | - |  |  |
| ***POLAND (n = 185)*** |  | *Species* | | | |  |  |  |  |  |
| Site | Size | COAU | NANA | NATE | ZALO |  |  |  |  |  |
| Kuyavia-Pomerania | 17969 | 2/0/12 | 0/0/15 | - | - |  |  |  |  |  |
| Lesser Poland | 15183 | 11/0/21 | 7/0/46 | - | - |  |  |  |  |  |
| Lower Silesia | 19947 | 0/0/2 | 1/0/1 | - | - |  |  |  |  |  |
| Opole | 9412 | 3/0/21 | 0/0/3 | - | - |  |  |  |  |  |
| Silesia | 12333 | 0/0/1 | - | 0/0/11 | - |  |  |  |  |  |
| Subcarpathia | 17846 | 0/0/1 | 4/4/10 | - | 3/1/41 |  |  |  |  |  |
| ***PORTUGAL (n = 54)*** |  | *Species* | | | | | |  |  |  |
| Site | Size | COAU | MAMO | NAAS | NAMA | VILA | VISE |  |  |  |
| Portalegre (Alentejo) | 6065 | - | - | - | 0/0/3 | - | - |  |  |  |
| Braga/Porto/Viana do Castelo (Norte) | 7323 | 1/0/4 | 0/0/1 | 0/0/5 | 0/0/1 | 4/0/34 | 0/0/6 |  |  |  |
| ***SPAIN (n = 155)*** |  | *Species* | | | | | | | | |
| Site | Size | COAU | COGI | MAMO | NAAS | NAMA | VIAS | VILA | VISE | ZASC |
| Huelva (Andalusia) | 3910 | - | 0/0/1 | - | - | - | - | - | - | 1/0/3 |
| Leon/Zamora/Burgos (Castile y Leon) | 40432 | 0/0/1 | 0/0/1 | - | 1/0/2 | 0/0/1 | 1/0/23 | 0/0/29 | 0/0/19 | - |
| Galicia | 29574 | 0/0/4 | - | - | 0/0/7 | 0/0/4 | - | - | 2/0/26 | - |
| La Rioja | 5045 | 0/0/1 | 0/0/1 | 0/0/1 | - | 0/0/2 | 1/0/11 | 0/0/12 | - | - |
| Navarra | 10391 | - | - | 0/0/1 | 0/0/1 | - | 0/0/4 | - | - | - |
| ***SWITZERLAND (n = 243)*** |  | *Species* | | | | | | | |  |
| Site | Size | COAU | HIVI | NAHE | NAMA | NATE | VIAS | VIBE | ZALO |  |
| Bern | 5960 | - | - | 1/0/6 | - | 14/15/26 | 0/0/2 | 0/0/3 | - |  |
| Neuchatel | 802 | - | - | 6/7/26 | - | - | - | 0/0/4 | - |  |
| Nidwalden | 276 | 0/0/2 | - | 0/0/1 | - | 6/5/7 | 0/0/2 | - | - |  |
| Obwalden | 491 | - | - | - | - | 5/6/10 | - | - | - |  |
| Schwyz | 908 | - | - | - | 1/1/1 | 0/0/1 | - | - | - |  |
| Ticino | 2812 | - | 2/3/11 | 4/3/15 | - | 3/4/15 | - | - | 2/1/4 |  |
| Vaud | 3212 | - | 3/1/30 | 19/19/55 | 0/0/2 | 11/0/16 | 0/0/4 | - | - |  |
| ***UKRAINE (n = 66)*** |  | *Species* | | | | | | | |  |
| Site | Size | COAU | DOCA | ELSA | NANA | NATE | VIBE | VINI | VIRE |  |
| Chernihiv | 31865 | - | - | - | - | - | 0/0/1 | - | - |  |
| Crimea | 26081 | - | - | 0/0/1 | - | - | - | - | 0/0/1 |  |
| Kharkiv | 31415 | 0/0/3 | - | - | 7/7/8 | 0/0/1 | - | 9/1/32 | 0/0/6 |  |
| Kherson | 28461 | - | 0/0/1 | - | 0/0/1 | - | - | - | - |  |
| Luhansk | 26684 | - | - | 0/0/2 | - | - | - | - | 0/0/5 |  |
| Mykolaiv | 24598 | - | 0/0/1 | - | - | - | - | - | - |  |
| Odesa | 33310 | 0/0/1 | 0/0/2 | - | - | - | - | - | - |  |

**Table S2.** Coefficients from Bayesian hierarchical model for species-level *Ophidiomyces ophidiicola* prevalence, the pathogen that causes snake fungal disease, in Europe (coefficient ± standard deviation (95% credible intervals)). The data were analyzed with a Bernoulli distribution and log link including country as a random effect and species as a population-level effect (*Coronella austriaca* (COAU), *Dolichophis caspius* (DOCA), *Hierophis viridiflavus* (HIVI), *Natrix astreptophora* (NAAS), *Natrix helvetica* (NAHE), *Natrix natrix* (NANA), *Natrix maura* (NAMA), *Natrix tessellata* (NATE), *Vipera ammodytes* (VIAM), *Vipera aspis* (VIAS), *Vipera berus* (VIBE), *Vipera latastei* (VILA), *Vipera nikolskii* (VINI), *Vipera renardi* (VIRE), *Vipera seoanei* (VISE), *Vipera ursini* (VIUR), *Zamenis longissimus* (ZALO)). The model is parameterized in relation to the reference level or intercept NATE, and each parameter represents the difference from this reference level.

|  | Estimate ± Standard Deviation (95% CI) |
| --- | --- |
| Intercept (NATE) | -1.85±0.51 (-3.02, -0.97) |
| Species COAU | -3.76±1.3 (-6.74, -1.73) |
| Species DOCA | -8.79±5.69 (-22.43, -1.12) |
| Species HIVI | -1.07±0.56 (-2.23, -0.02) |
| Species NAAS | -8.5±5.8 (-21.77, -0.39) |
| Species NAHE | -0.16±0.38 (-0.9, 0.58) |
| Species NAMA | -2.66±1.31 (-5.7, -0.56) |
| Species NANA | -0.75±0.49 (-1.73, 0.23) |
| Species VIAM | -8.04±6.17 (-22.98, 0.2) |
| Species VIAS | -9.44±5.25 (-22.15, -2.54) |
| Species VIBE | -3.09±1.28 (-6.13, -1.11) |
| Species VILA | -8.97±5.11 (-21.12, -1.92) |
| Species VINI | -5±1.38 (-8.06, -2.65) |
| Species VIRE | -10.25±5.57 (-23.46, -2.78) |
| Species VISE | -8.87±5.33 (-21.25, -1.43) |
| Species VIUR | -8.39±5.84 (-22.5, -0.12) |
| Species ZALO | -0.94±0.51 (-1.98, 0.07) |

**Table S3.** Coefficients from Bayesian hierarchical model for differences in severity of snake fungal disease (SFD) among species in Europe for individuals that were confirmed positive for the pathogen that causes SFD, *Ophidiomyces ophidiicola,* through quantitative polymerase chain reaction (qPCR) (coefficient ± standard deviation (95% credible intervals)). The data were analyzed with a binomial distribution that included site and individual snake ID as random effects, and species as a population-level effect. The model is parameterized in relation to the reference level or intercept *Natrix tessellata* and each parameter represents the difference from this reference level.

|  | Estimate ± Standard Deviation (95% CI) |
| --- | --- |
| Intercept *N. tessellata* | -6.08±0.22 (-6.5, -5.65) |
| species *C.* austriaca | 1.08±1.12 (-1.14, 3.21) |
| species *H.* viridiflavus | 0.11±0.59 (-1.06, 1.22) |
| species *N.* helvetica | -0.47±0.37 (-1.19, 0.28) |
| species *N.* maura | 0.72±1.11 (-1.48, 2.91) |
| species *N.* natrix | 0.73±0.44 (-0.14, 1.58) |
| species *V.* berus | -0.13±9.92 (-19.53, 19.46) |
| species *V. nikolskii* | -0.07±10.28 (-19.96, 20.25) |
| species *Z. longissimus* | 0±0.48 (-0.94, 0.96) |

**Table S4.** Alignment of representative sequences previously deposited in GenBank that correspond to the four internal transcribed spacer 2 (ITS2) region genotypes detected in our study.

```
Genotype_IB_KY474061	GAAATGCGATAAGTAATGTGAATTGCAGAATTCCGTGAATCATCGAATCTTTGAACGCAC     60
Genotype_IIF_KX148658.1	GAAATGCGATAAGTAATGTGAATTGCAGAATTCCGTGAATCATCGAATCTTTGAACGCAC     60
Genotype_IA_KY474059.1	GAAATGCGATAAGTAATGTGAATTGCAGAATTCCGTGAATCATCGAATCTTTGAACGCAC     60
Genotype_IIDE_OL457490.1	GAAATGCGATAAGTAATGTGAATTGCAGAATTCCGTGAATCATCGAATCTTTGAACGCAC     60
			************************************************************

Genotype_IB_KY474061	ATTGCGCCCCCTGGTATTCCGGGGGGCATGCCTGTCCGAGCGTCATTGCAACCCCCTCAA     120
Genotype_IIF_KX148658.1	ATTGCGCCCCCTGGTATTCCGGGGGGCATGCCTGTCCGAGCGTCATTGCAACCCCCTCAA     120
Genotype_IA_KY474059.1	ATTGCGCCCCCTGGTATTCCGGGGGGCATGCCTGTCCGAGCGTCATTGCAACCCCCTCAA     120
Genotype_IIDE_OL457490.1	ATTGCGCCCCCTGGTATTCCGGGGGGCATGCCTGTCCGAGCGTCATTGCAACCCCCTCAA     120
			************************************************************

Genotype_IB_KY474061	GCCCGGCTTGTGTGTTGGGGGCGCCCGCCCCGAAGTCCTCGGGCGCGGGCCCCCCCCCAA     180
Genotype_IIF_KX148658.1	GCCCGGCTTGTGTGTTGGGGGCGCCCACCCCGAAGTCCTCGGGCGCGGGCCCCCCCCCAA     180
Genotype_IA_KY474059.1	GCCCGGCTTGTGTGTTGGGGGCGCCCGCCCCGAAGTCCTCGGGCGCGGGCCC-CCCCCAA     179
Genotype_IIDE_OL457490.1	GCCCGGCTTGTGTGTTGGGGGTGCCCACCCCGAAGTCCTCGGGCGCGGGCCC-CCCCCAA     179
			********************* **** ************************* *******

Genotype_IB_KY474061	ATGCAGTGGCGGCACCGAGTTCCTGGTGTCTGAGTGTATGGGAATCTGTTTCTGTCTCGC     240
Genotype_IIF_KX148658.1	ATGCAGTGGCGGCACCGAGTTCCTGGTGTCTGAGTGTATGGGAATCTGTTTCTGTCTCGC     240
Genotype_IA_KY474059.1	ATGCAGTGGCGGCACCGAGTTCCTGGTGTCTGAGTGTATGGGAATCTGTTTCTGTCTCGC     239
Genotype_IIDE_OL457490.1	ATGCAGTGGCGGCACCGAGTTCCTGGTGTCTGAGTGTATGGGAATCTGTTTCTGTCTCGC     239
			************************************************************

Genotype_IB_KY474061	TCGAAGACCCGATCGGCGCCCGTCGTCAACCCCC     274
Genotype_IIF_KX148658.1	TCGAAGACCCGATCGGCGCCCGTCGTCAACCCCC     274
Genotype_IA_KY474059.1	TCGAAGACCCGATCGGCGCCCGTCGTCAACCCCC     273
Genotype_IIDE_OL457490.1	TCGAAGACCCGATCGGCGCCCGTCGTCAACCCCC     273
			**********************************
```

**Table S5.** Bayesian model comparisons explaining *Ophidiomyces ophidiicola* prevalence across the landscape using the leave-one-out cross-validation (LOO).

| **Model comparison tested** | | |  |  |  |  |  |  |
| --- | --- | --- | --- | --- | --- | --- | --- | --- |
|  | elpd_diff | se_diff | elpd_loo | se_elpd_loo | p_loo | se_p_loo | looic | se_looic |
| clade.detection*Species+(1\|Site) | 0 | 0 | -265.349 | 19.198 | 27.025 | 4.366 | 530.697 | 38.396 |
| clade.detection+Species+(1\|Site) | -8.705 | 5.991 | -274.053 | 18.622 | 23.138 | 3.641 | 548.107 | 37.244 |
| Species+(1\|Site) | -10.086 | 6.059 | -275.434 | 19.110 | 23.835 | 3.949 | 550.869 | 38.221 |
| clade.detection+(1\|Site) | -23.388 | 10.734 | -288.737 | 17.205 | 13.931 | 1.312 | 577.473 | 34.410 |

**Table S6.** Coefficients from Bayesian hierarchical model of interaction between species and clade explaining *Ophidiomyces ophidiicola* prevalence in Europe (coefficient ± standard deviation (95% credible intervals)). The data were analyzed with a binomial distribution which included site as a random effect, and species and clade as population-level effects. The model is parameterized in relation to the reference level or intercept *Natrix tessellata* and each parameter represents the difference from this reference level.

|  | Estimate ± Standard Deviation (95% CI) |
| --- | --- |
| Intercept Clade Europe: *N. tessellata* | -0.27±0.68 (-1.56,1.13) |
| Clade North_America: *N. tessellata* | -0.61±0.89 (-2.48,1.09) |
| Species *N. helvetica* | -1.2±1.11 (-3.52,0.85) |
| Species *N. natrix* | -1.82±0.66 (-3.16,-0.59) |
| Species *Z. longissimus* | -3.92±1.1 (-6.22,-1.88) |
| Clade North_America:species *N. helvetica* | 1.25±1.2 (-0.97,3.74) |
| Clade North_America:species *N. natrix* | 2.93±1.08 (0.88,5.11) |
| Clade North_America:species *Z. longissimus* | 4.12±1.3 (1.69,6.8) |

**Table S7.** Coefficients from Bayesian hierarchical model of interaction between species and clade explaining *Ophidiomyces ophidiicola* prevalence in Europe (coefficient ± standard deviation (95% credible intervals)) with spatial conditional autoregressive term. The data were analyzed with a binomial distribution which included species and clade as population-level effects, and an adjacency matrix and grouping factor of site. The model is parameterized in relation to the reference level or intercept *Natrix tessellata* and each parameter represents the difference from this reference level.

|  | Estimate ± Standard Deviation (95% CI) |
| --- | --- |
| Intercept Clade Europe: *N. tessellata* | -0.14±0.45 (-1.02,0.70) |
| Clade North_America: *N. tessellata* | -0.59±0.54 (-1.66,0.42) |
| Species *N. helvetica* | -1.21±0.69 (-2.49,0.08) |
| Species *N. natrix* | -1.92±0.60 (-3.01,-0.76) |
| Species *Z. longissimus* | -4.36±1.22 (-7.35,-2.06) |
| Clade North_America:species *N. helvetica* | 0.61±0.74 (-0.85,1.95) |
| Clade North_America:species *N. natrix* | 1.30±1.02 (-0.63,3.32) |
| Clade North_America:species *Z. longissimus* | 3.14±1.31 (0.63,6.24) |

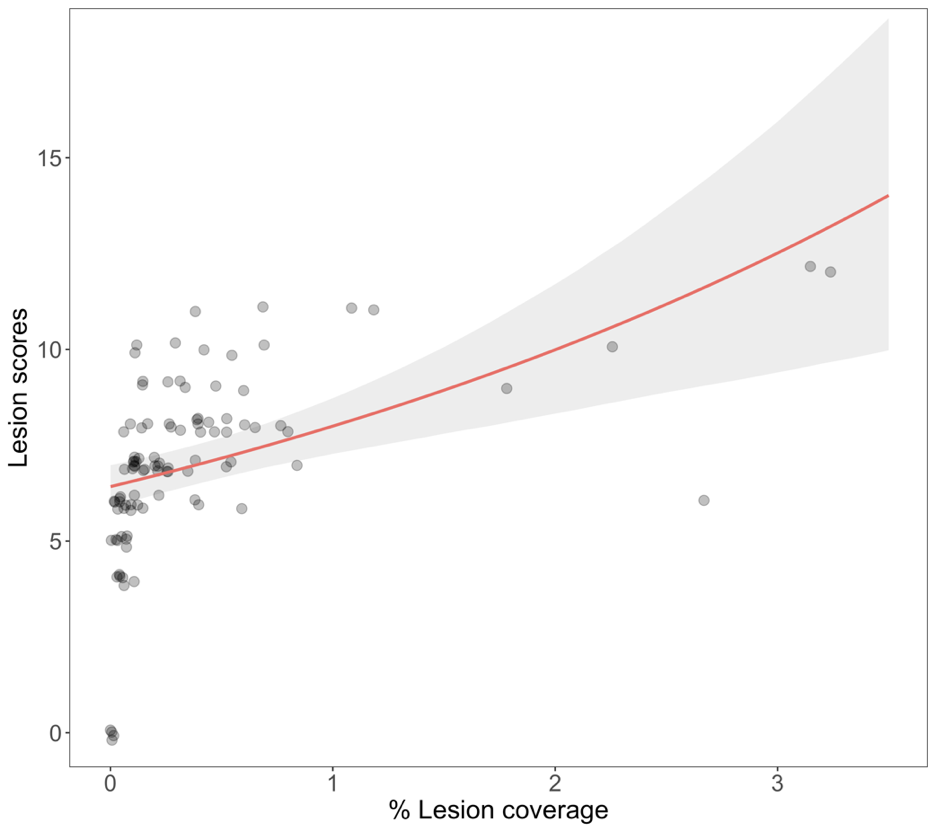

**Figure S1.** Relationship between lesion scores and lesion coverage (percentage of the snake’s total surface area covered in lesion) measured from photos.

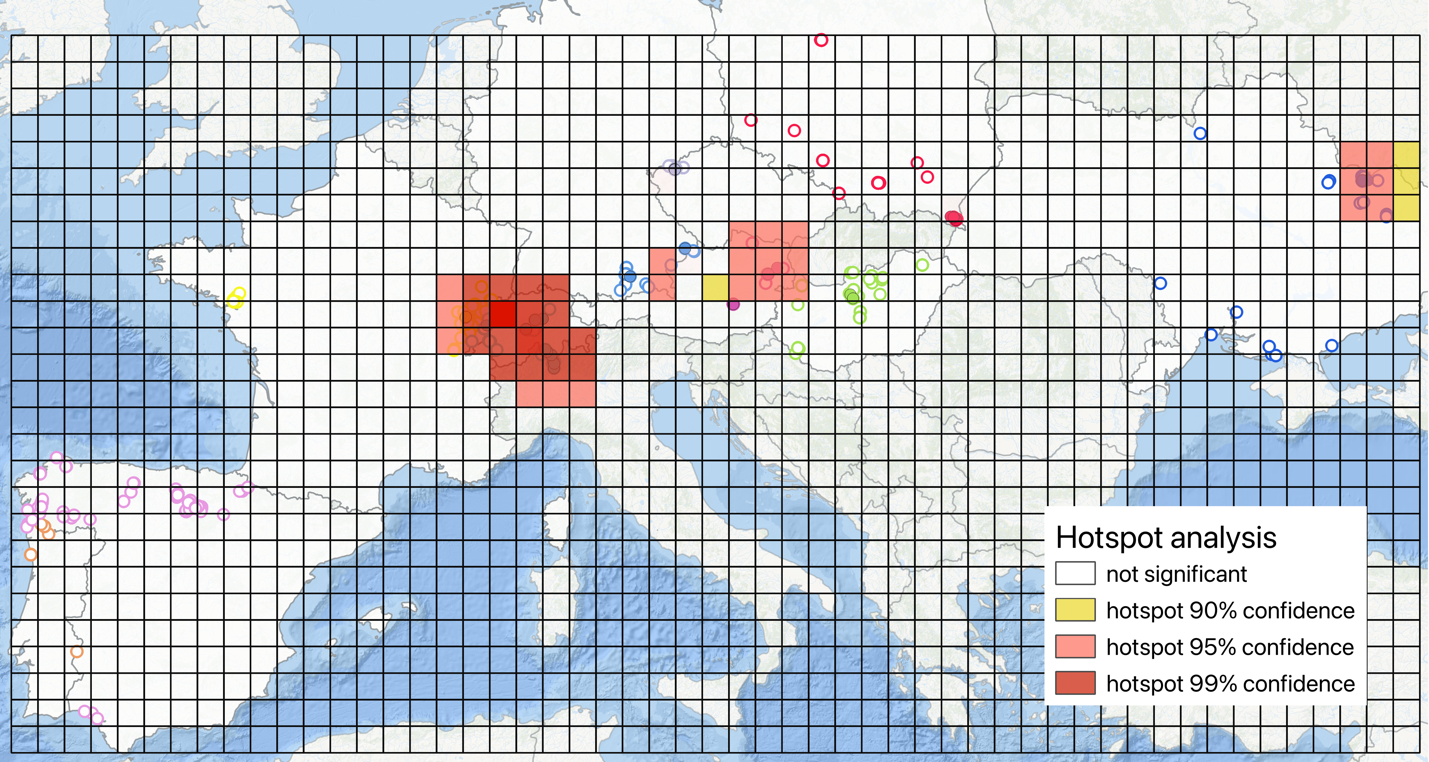

**Figure S2.** Hotspot analysis following Getis-Ord Gi^*^ method with Local Indicators of Spatial Association (LISA) statistics.

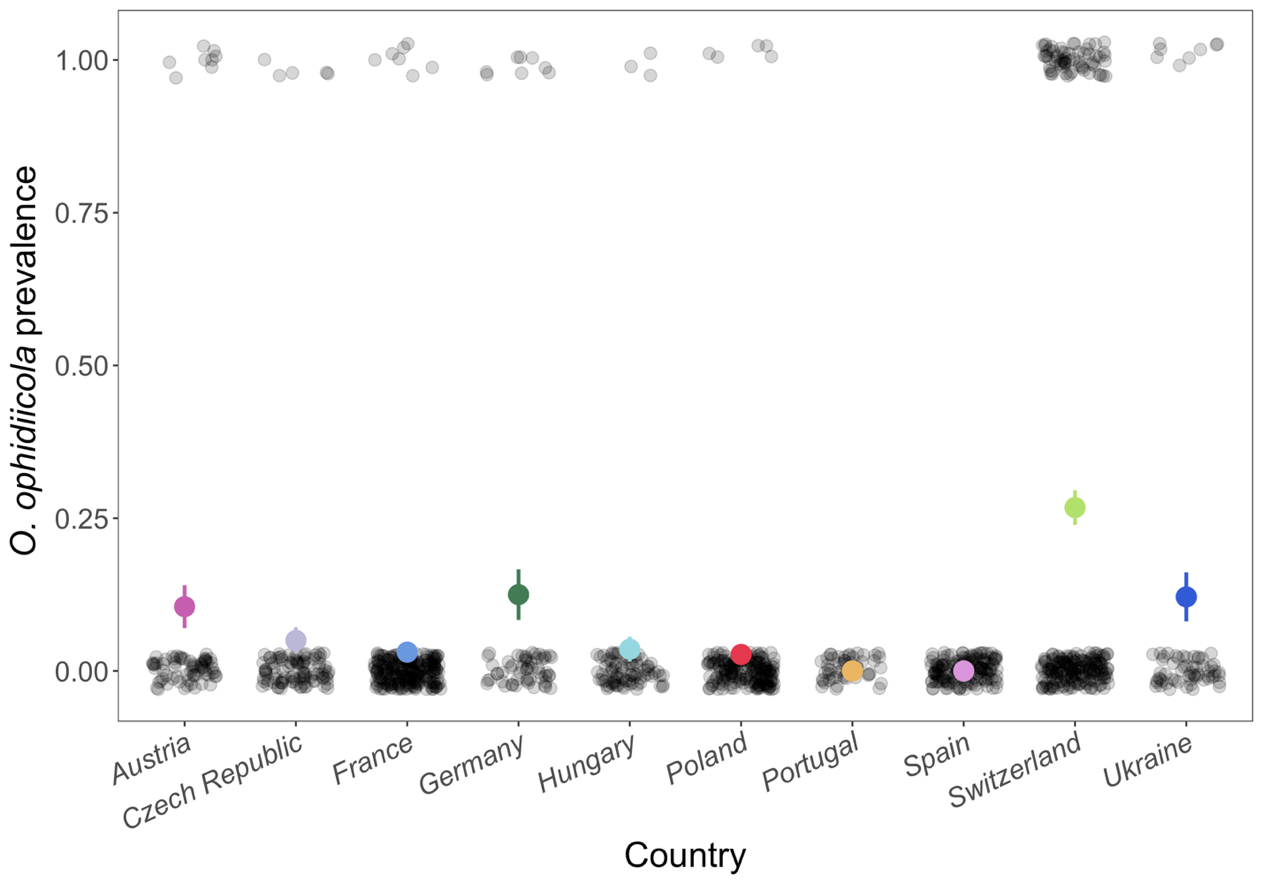

**Figure S3.** Observed *Ophidiomyces ophidiicola* prevalence across all countries sampled using quantitative polymerase chain reaction (qPCR). Each black point represents a single snake as being either negative (prevalence = 0) or positive (prevalence = 1). Color circles are mean prevalence ± standard deviation for each country.

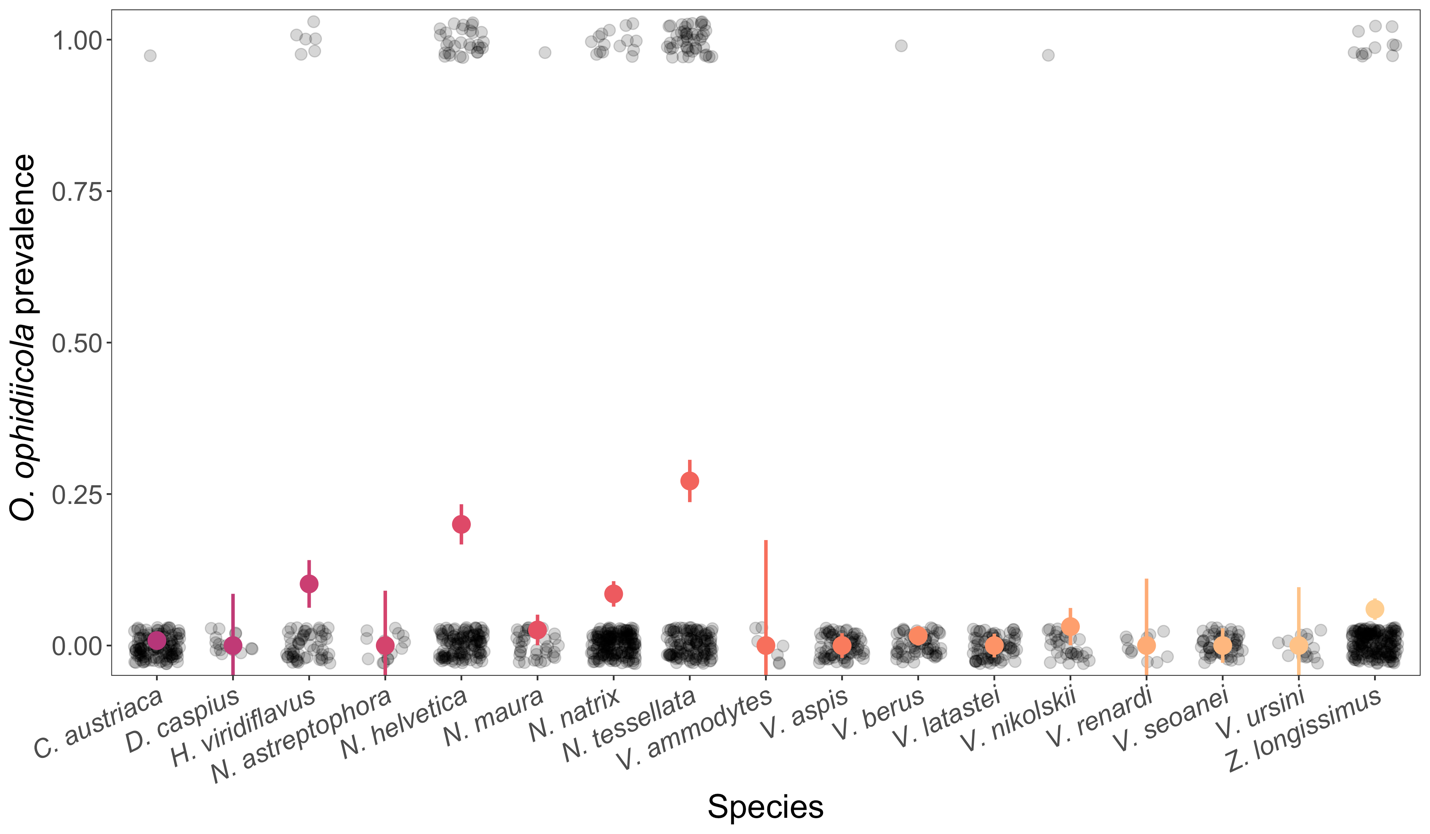

**Figure S4.** Observed *Ophidiomyces ophidiicola* prevalence across all species sampled using quantitative polymerase chain reaction (qPCR). Each black point represents a single snake as being either negative (prevalence = 0) or positive (prevalence = 1). Color circles are mean prevalence ± standard deviation for each species.

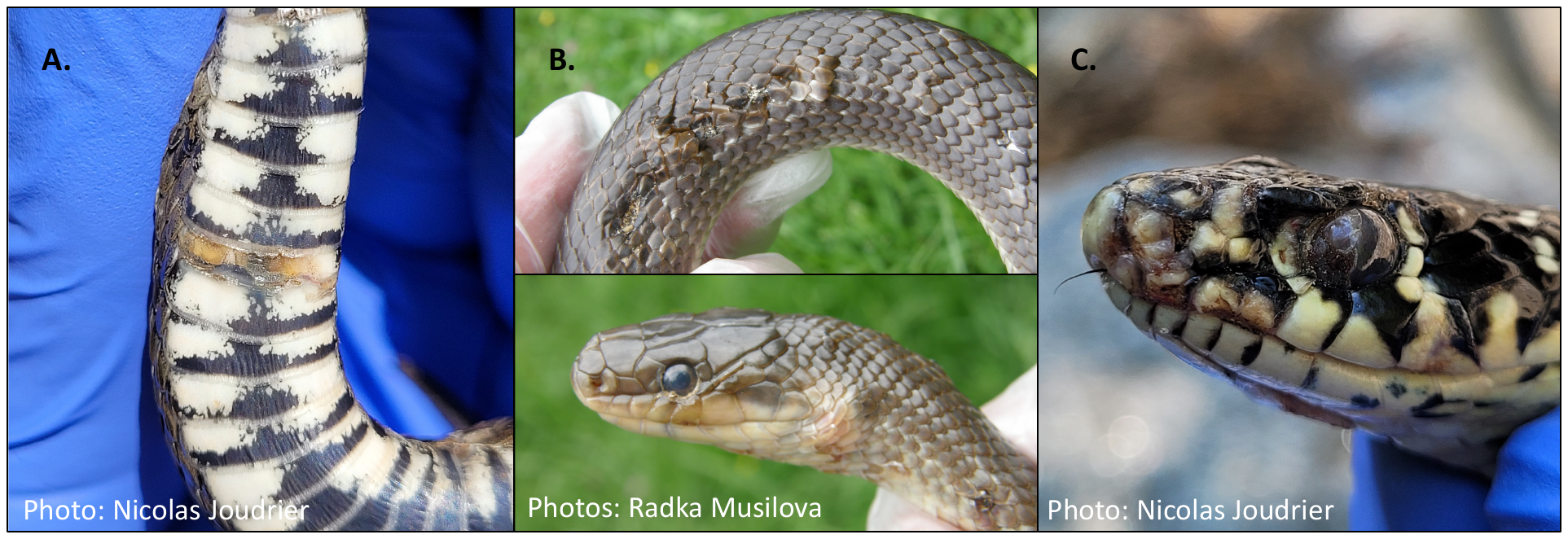

**Figure S5.** Photos of lesions from snakes that were qPCR positive for *Ophidiomyces ophidiicola*. (A) Photo of a mild ventral lesion on *Natrix tessellata* in Switzerland. (B) Photo of moderate lesions on the head and body of *Zamenis longissimus* in Czech Republic. (C) Photo of a severe lesion on the head of *Hierophis viridiflavus* in Switzerland.

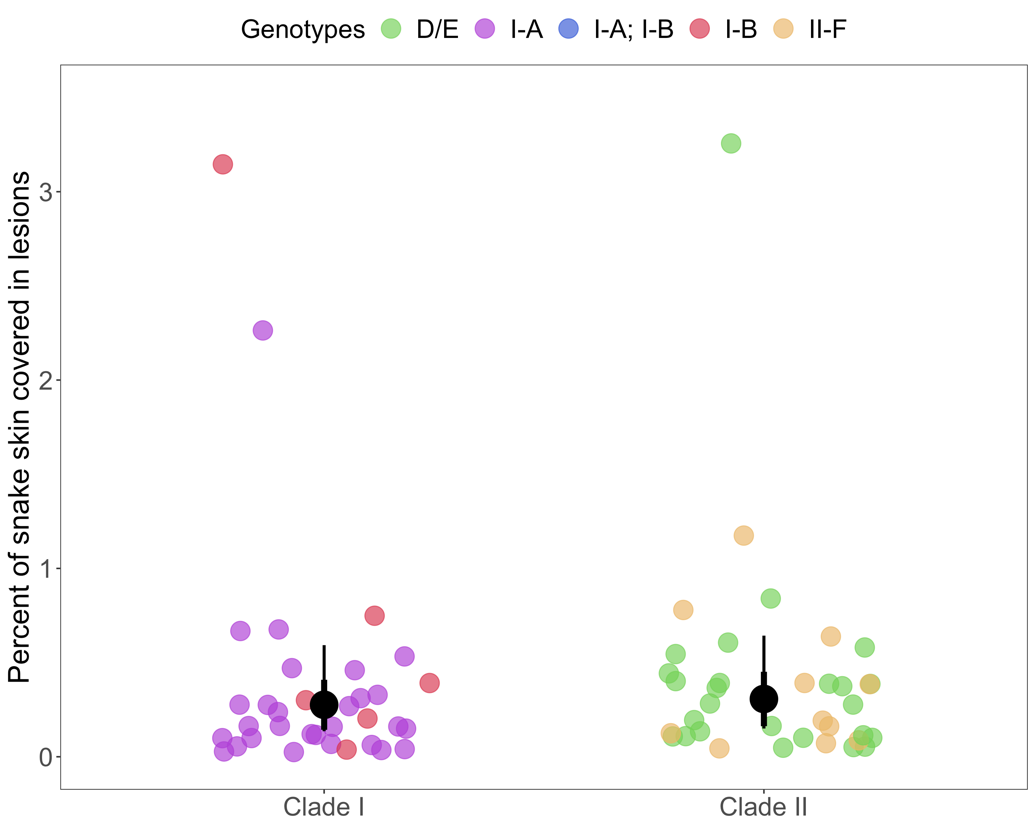

**Figure S6.** Variation in disease severity (fraction of snake covered in lesions) by pathogen clade infecting the host. Each point is a single snake for which *Ophidiomyces ophidiicola* clade was determined, colors represent the different lineages of *O. ophidiicola*. The black circles and whiskers show the model predicted posterior mean, ± standard deviation (thick lines), and 95% credible intervals (thin lines).
